# An integrative approach for profiling antibody responses in bats to human pathogens

**DOI:** 10.1101/2024.11.29.626104

**Authors:** Nia Toshkova, Violeta Zhelyzkova, Kaloyana Koseva, Katrin Dimitrova, Farida Elshaer, Robin V. Lacombe, Maxime Lecerf, Anastas Pashov, Jordan D. Dimitrov

## Abstract

Serological analyses are a fundamental tool for identifying infections by a wide range of pathogens. They offer a current overview of pathogen prevalence and insight into past infections. This is particularly relevant for bats, given their high capacity to tolerate pathogens and their role as reservoirs of zoonotic diseases. At present, serological studies in bats have predominantly employed traditional techniques such as enzyme-linked immunosorbent assay (ELISA). However, these techniques have several limitations, including low throughput and the lack of bat-specific detection antibodies. To address these limitations, we developed an integrative approach for systemic serological analyses based on microarray technology, which enables the simultaneous detection of bat IgG antibodies against >190 human pathogens (viruses, bacteria, protists). The results of our analyses demonstrated an antibody response in bats targeting multiple epitopes from different pathogens, thereby proving the method’s high-throughput capability. Furthermore, this approach does not rely on the use of IgG detection reagents, thereby allowing for its application to a diverse range of bat species. This assay offers insights into the infections of bats with pathogens, thereby enhancing our comprehension of zoonotic disease dynamics and facilitating targeted pathogen surveillance.

## Introduction

Bats (order *Chiroptera*) are the only mammals that have evolved the capacity for sustained flight, which is associated with some unique physiological traits. Bats exhibit unusually long lifespans, resistance to age-related diseases, and an outstanding potential to tolerate viral infections (1, 2). The latter is attributed to genetic adaptations in innate and adaptive immune components, which results in both heightened pathogen resistance and dampened inflammatory responses (2–6). The elevated pathogen tolerance, the number and diversity of existing species, and certain ecological traits (e.g. roosting aggregation) are considered to be key factors contributing to the major role of bats as a reservoir of zoonotic diseases (7–12). Indeed, bats have been documented to harbor a considerable diversity of viruses, including pathogens such as Ebola, Marburg, Lassa, Nipah, Hendra, and Rabies viruses, as well as closely related viruses to SARS-CoV-1, MERS, and SARS-CoV-2(2, 7, 9). Additionally, many pathogenic bacteria and protists have been detected in bats (13).

The high potential for the emergence and spillover of zoonotic diseases necessitates a systematic approach to monitoring pathogen dynamics in bat populations. (7). Serological methods are well-suited to this objective, as fluctuations in pathogen-specific antibody levels in blood can provide insights into past and ongoing infections. In this context, classical enzyme-linked immunosorbent assay (ELISA)-based techniques and multiplexed immunoassay (Luminex) have demonstrated efficacy in detecting antibodies to diverse human pathogens in bats (14–18). These assays, however, have significant limitations. They are relatively low throughput and are most often used for the detection of antibodies to predefined pathogens. Another challenge in bat serological research is the limited availability of reagents for the detection of bat immune molecules. Although bats are the second most diverse mammalian order with more than 1400 different species, such reagents are available only for a few of them, emphasizing the need for further development in this area.

To address these limitations, a novel approach, Phage ImmunoPrecipitation Sequencing (PhIP-Seq VirScan), has recently been applied to bat samples with the objective of comprehensively profiling their exposure history to the human virome (19, 20). Despite its effectiveness, this method is complex, focused only on viruses, and requires a greater investment of time compared to standard serological assays.

In this study, we propose an integrative strategy that enables unbiased, comprehensive evaluation of bat antibody specificity against a vast array of human pathogens, encompassing bacteria, viruses, and protists. This approach can be completed within a single day, does not rely on antibody detection reagents, and can be applied to multiple bat species. Implementing this assay offers an effective approach for monitoring pathogen dynamics in bat populations and identifying priority host species and pathogens for targeted zoonotic surveillance.

## Results and Discussion

### Optimization of the assay

The goal of this study was to develop an assay that enables rapid, untargeted monitoring of bat IgG specificities towards a broad repertoire of human pathogens. Furthermore, our objective was to eliminate the requirement for specific antibody detection reagents, thereby enabling the method to be applied to a diverse range of bat species. The methodology, illustrated in **Fig. 1**, comprises three steps: 1) antibody purification from pooled bat sera (n=35 individuals) using IgG purification reagents with broad specificity among mammals; 2) direct antibody labeling with high-quantum yield fluorochromes; and 3) high-throughput evaluation of bat IgG immunoreactivity using a pathogen microarray platform.

**Figure 1.**
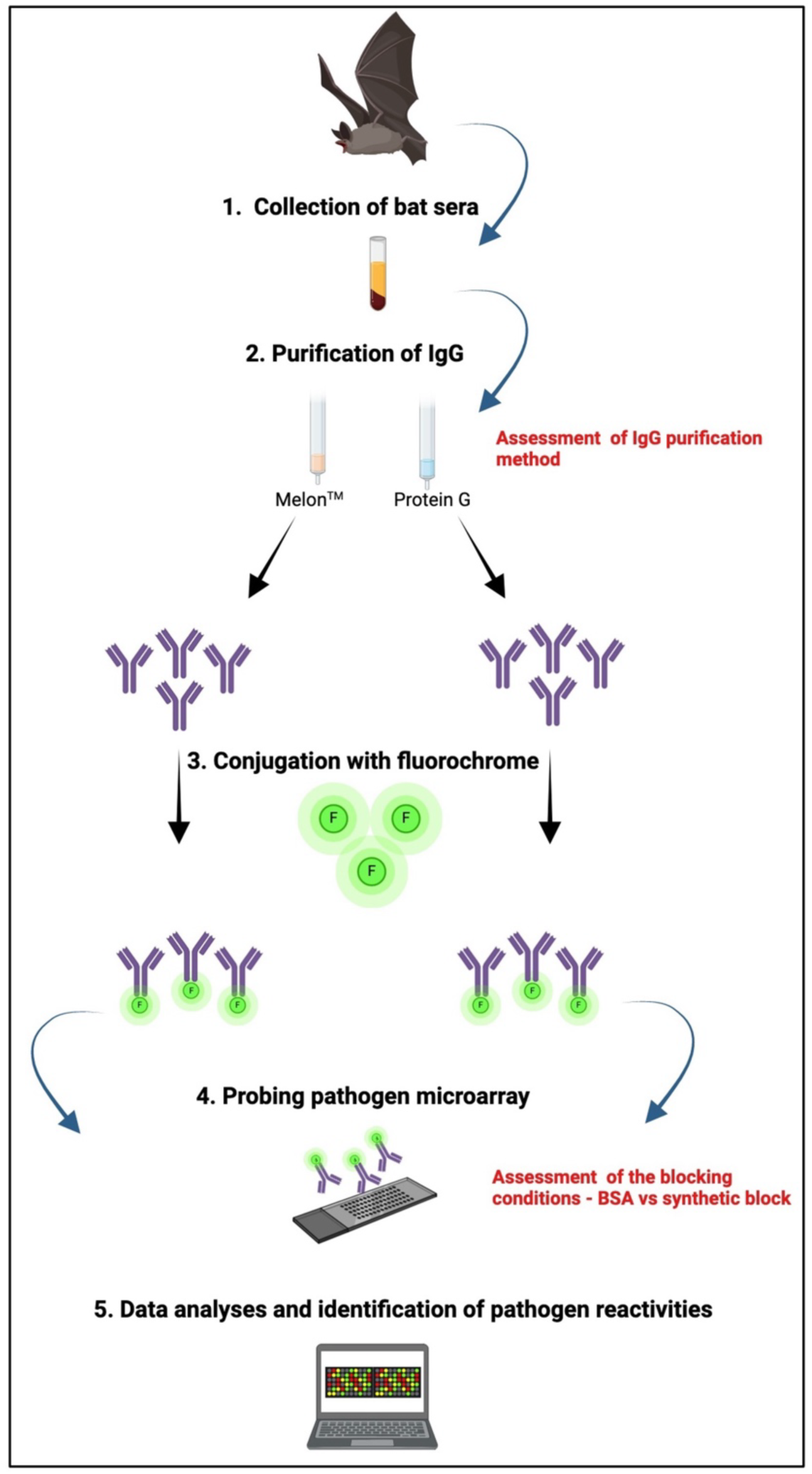
Schematic representation of the experimental approach. The optimization steps are highlighted in red. The scheme was created by using BioRender (BioRender.com).

To develop and optimize the approach, sera samples from the common noctule bat (*Nyctalus noctula*, Schreber, 1774) were usde as a model. In order to purify IgG, two reagents with broad efficacy across different orders of mammals were compared: Protein G and Melon™ (Thermo Fisher Scientific). Protein G is a specific binder of IgG, requiring exposure to a low pH (<3) for the elution of bound antibodies. In contrast, Melon^TM^ is based on ion-exchange chromatography, wherein all serum proteins, except IgG, remain bound to the matrix. Consequently, it purifies antibodies in mild conditions (pH 6.5-7). Our data demonstrated that both Protein G and MelonTM successfully purified IgG from pooled bat *N. noctula* sera with high quality (**Suppl. Fig. 1**).

Subsequently, the purified bat IgG was labeled with Alexa Fluor^™^ 555 NHS ester, which forms a covalent amide bond with the primary amino groups of proteins. This step allows for the assessment of IgG immunoreactivity without the need for secondary antibodies or other IgG detection reagents. The labeling procedure is rapid, taking less than two hours, and can be readily adapted for high-throughput applications. The final step of the assay is dedicated to profiling the pathogen reactivities of bat IgG to a broad repertoire of antigens representing a variety of human pathogens. To this end, we used a high-density peptide array, namely the PEPperCHIP^®^ Infectious Disease Epitope Microarray (PEPperPRINT), which was originally developed for the purpose of serological profiling of infections in humans. This array displays over 3,700 linear B cell epitopes derived from proteins of 196 human pathogens, including viruses, bacteria, and protists. It should be noted that this array has only been selected as a proof of concept. Alternative antigen arrays, either standard or customized, can be employed to address specific objectives.

The experiment was alternatively performed by using two blocking agents for microarrays: a solution of BSA and a solution of synthetic polymers (a synthetic block, Thermo Fisher Scientific). The antibody binding analyses demonstrated that the IgG purification method had a significant impact on the overall reactivity of bat antibodies. Thus, the antibodies purified by Protein G exhibited higher binding intensity (MFI) and binding to a greater number of pathogen-derived epitopes in comparison to the antibodies purified under mild conditions (p < 0.0001, non-parametric Wilcoxon test) (**Fig. 2a, b**). These discrepancies were substantiated by Z-score calculation (**Fig. 2c**) and hierarchical clustering (**Fig. 2d**). Conversely, the type of blocking agent employed did not markedly influence the reactivity of bat IgG to pathogen-derived epitopes (**Fig. 2**).

**Figure 2.**
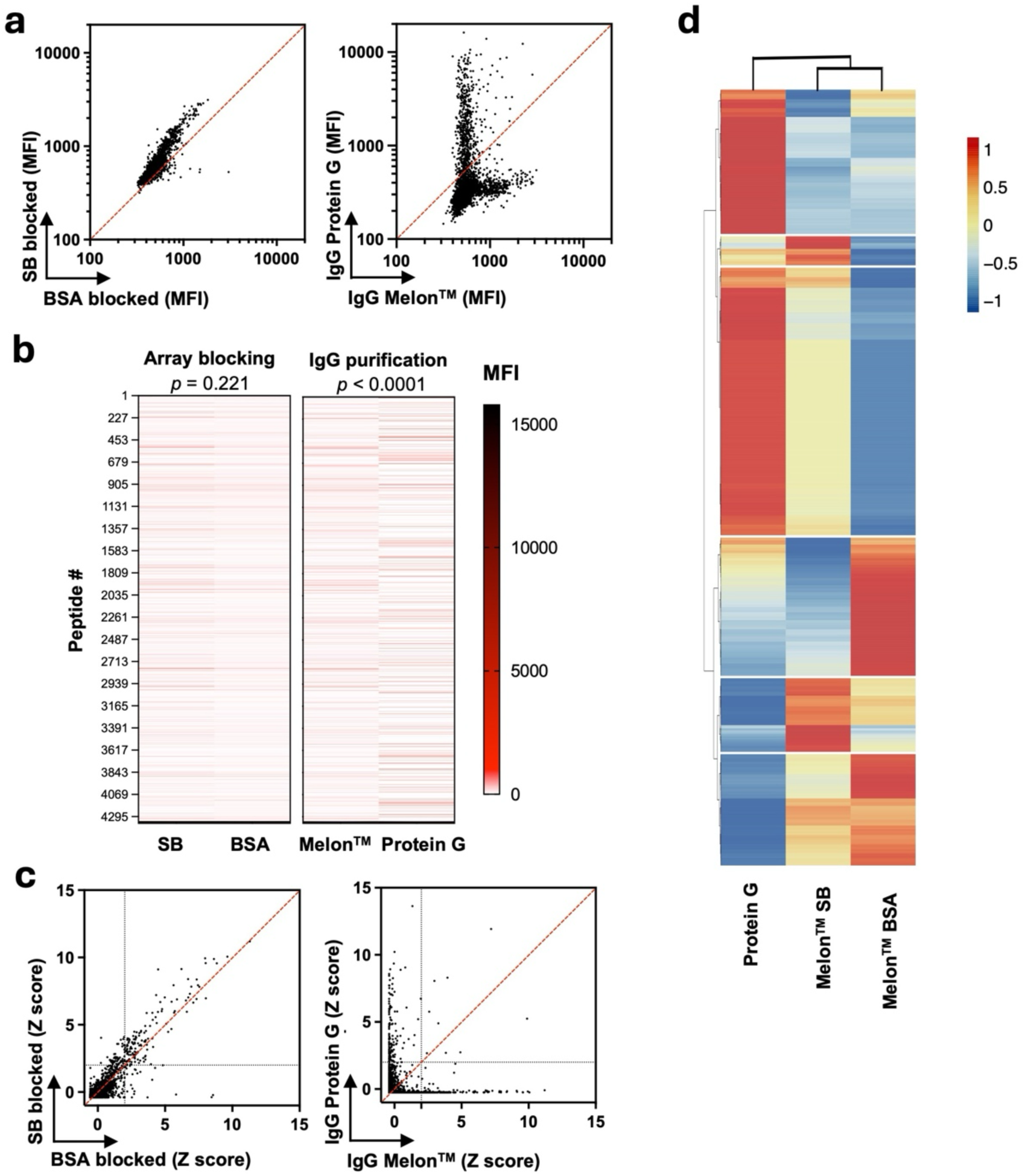
High throughput analyses of reactivity of bat antibodies. **a**. Plots depicting the mean fluorescence intensity of the raw signal from microarray chips. Each spot indicates the binding intensity of bat antibodies to a single peptide. **b**. Heat-map of MFI values obtained after subtraction of the background. Each line in the graph represents the reactivity to a single peptide in the array. Statistical analyses were performed by a Wilcoxon two-tailed test to compare the mean antibody binding reactivity for the indicated conditions. **c**. Plots depicting the values of Z scores. The dashed gray lines correspond to Z score of 2, which is considered as a threshold of significance. Each dot in the plots corresponds to the Z score of antibody reactivity to a single peptide. **d**. Heat map and hierarchical clustering of Z scores depicting the reactivity of bat antibodies to peptides in microarrays. Each column in the heat maps displays the binding intensity of a single peptide. The average of each column in the map is always equal to 0. The map distinguished six clusters of peptides.

The observed disparity in antigen reactivity of bat IgG as a function of the purification methodology can be attributed to the step involving exposure to low pH in the case of Protein G. Low pH values, as utilized for the elution of antibodies from Protein G, have the potential to induce irreversible alterations in a subset of IgG molecules (21, 22). Moreover, previous studies have demonstrated that transient exposure to pH<4 can induce antigen-binding polyreactivity in both monoclonal and polyclonal human and mouse IgG (23, 24). The induction of polyreactivity by acidic elution from Protein G provides a plausible explanation for the overall increased reactivity of bat IgG to pathogen-derived peptides. Based on these data, it is recommended that mild purification conditions be used for the isolation of bat IgG. Furthermore, the experimental time using the Melon^™^ system is considerably shorter, approximately 10 minutes, as compared to one using Protein G, which requires approximately 40 minutes.

All binding assays conducted in the present study were performed at standard room temperature (22-24 °C). Recent research by our laboratory has demonstrated that bat antibodies (including IgG from *N. noctula*) exhibit enhanced antigen binding diversity at elevated temperatures (>40 °C), which are typical for the active metabolic states of bats during flight (Toshkova, 2024 #1). Consequently, further experiments should assess the role of temperature in pathogen identification via the proposed approach.

### Identification of pathogen specificities of bat IgG

In order to ascertain the pathogen-specificities of bat antibodies, Z-scores were calculated for each epitope on the array, utilizing the values of mean fluorescence intensities. A Z-score value of ≧2 (p < 0.05) was deemed indicative of a positive hit. The statistical analyses demonstrated a notable reactivity of bat antibodies to a range of human viruses, bacteria, and protists (**Fig. 3 a,b**). Of particular interest is the observation of antibody binding to certain pathogens, including *Borrelia burgdorferi*, *Mycobacterium tuberculosis*, *Porphyromonas gingivalis*, *Plasmodium falciparum*, *Toxoplasma gondii*, *Trypanosoma cruzi*, and SARS-CoV, which exhibited reactivity across multiple epitopes belonging to diverse pathogen proteins. (**Fig. 3a**). For example, the bat antibodies in question exhibited specific recognition of many epitopes derived from nine, six, four, and two distinct proteins associated with *M. tuberculosis*, *P. falciparum*, *B. burgdorferi*, and SARS CoV, respectively (**Fig. 3a**). The observed broad specificity of the antibody response to multiple epitopes present in diverse proteins indicates that the analyzed *N. noctula* bats have mounted specific humoral immune responses to closely related pathogens present in bats. Conversely, in the case of certain pathogens (Puumala virus, Influenza B virus, HIV-1, Andes virus, etc.), bat antibodies demonstrated recognition of a single epitope derived from a single protein. It is likely that these results are due to non-specific antibody reactivity to peptides, rather than indicating specific immune responses. It is noteworthy that while IgG purification methods affected microarray immunofluorescence signals, the core set of identified pathogens (those with the highest Z-scores) remained consistent in both cases (**Fig. 3 a,b**). This corroborates the reliability of our findings and justifies the utilization of Z-score thresholds for the analysis of extensive immunoreactivity datasets, thereby facilitating the elimination of non-specific effects. The purification of IgG by Protein G resulted in a broader set of recognized antigens, but its effects were most often limited to significant binding (Z score ≧ 2) to a single epitope of a given pathogen, which was probably due to non-specific effects. It is worth noting that in the case of antibody binding to antigens of *Borrelia burgdorferi* and *Trypanosoma cruzi*, purification with Protein G resulted in a reduction in the number of significantly recognized epitopes in contrast to IgG purified by ion-exchange matrix (**Fig. 3 a,b**).

**Figure 3.**
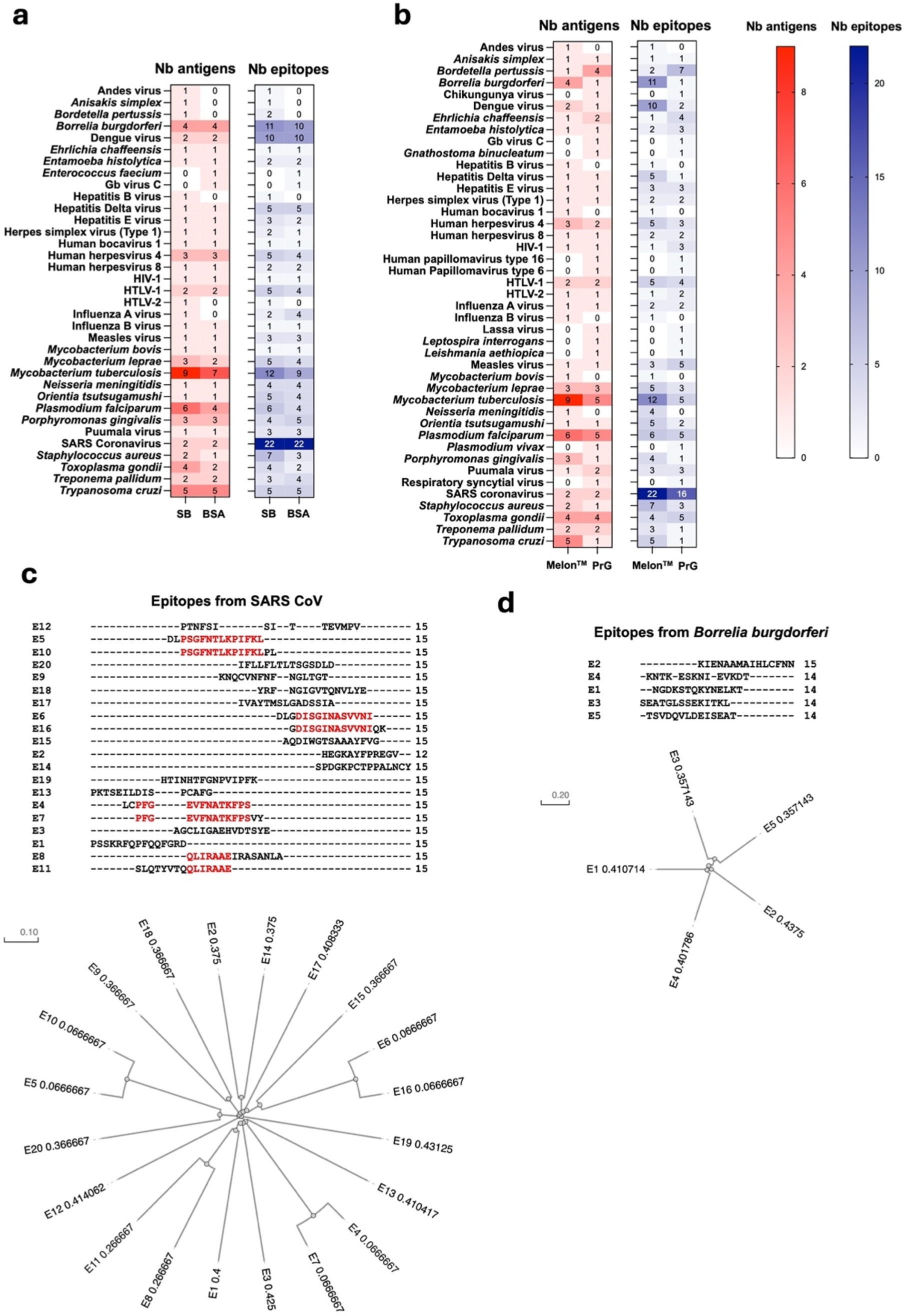
Identification and validation of specificities of bat antibodies for human pathogens. **a.** and **b**. Heat maps indicating the number of pathogen-derived proteins (in red) and the number of epitopes (in blue) recognized by bat antibodies with Z score ≧2. The heat maps compare the effect of the blocking agent (a) or the effect of IgG purification procedure (b). **c.** and **d.** Sequence alignment of epitopes from SARS CoV E2 glycoprotein precursor (c) and hypothetical lipoprotein from *Borrelia burgdorferi* (d) that are recognized with Z scores 2 or greater. The red color is used to highlight the identical motives. The sequence alignment plot and phylogenetic analyses (Guide trees) were generated by the Clustal Omega program (version 1.2.4) for multiple sequence alignment (https://www.ebi.ac.uk/).

Bat IgG antibodies were observed to recognize multiple epitopes originating from a single pathogen protein (**Fig. 3 a,b**). For example, 20 distinct epitope recognitions were identified for a single protein from SARS-CoV, specifically the E2 glycoprotein precursor. Similarly, five epitopes from a hypothetical lipoprotein of *Borrelia burgdorferi* were found to be specifically bound by bat IgG. To determine whether the multitude of epitopes from a single antigen reflects the recognition of a common motif, sequence alignment analyses were conducted (**Fig. 3c, d** and **Suppl. Fig. 2**). These analyses demonstrated that the majority of epitopes identified as positive hits lack a common sequence despite originating from the same antigen. For example, among the 20 epitopes from the E2 glycoprotein precursor that were recognized by bat antibodies, only four peptide pairs shared a common sequence motif. These data indicate that the antibodies specifically recognize 16 unique epitopes in the viral protein. Similarly, high diversity was also detected in the case of epitopes present in the hypothetical lipoprotein of *Borrelia burgdorferi* (**Fig. 3d**) as well as in proteins from other pathogens (**Suppl. Fig. 2**).

Taken together the data provide compelling evidence that the detected antibody specificities are indicative of specific immune recognition and not a consequence of non-specific cross-reactivities. This lends further support to the reliability of the microarray platform.

To further validate the capacity of the proposed approach to detect specific bat IgG antibodies, we applied a standard ELISA immunoassay with entire antigens from selected pathogens and with whole inactivated *M. tuberculosis* (**Fig. 4**). This bacterium was selected as one of the most prominently recognized pathogens by IgG of *N. noctula* bat in the microarray. In these experiments, the bat IgG antibodies were tested without fluorochrome labeling, and their binding to antigens was detected using biotinylated Protein G. This setting also controlled for potential artifacts due to the conjugation of IgG molecules with fluorochromes. The data obtained confirmed the pathogen reactivities identified by the microarray platform (**Fig. 4**). Furthermore, it confirmed that the IgG antibodies purified by Protein G displayed a tendency for higher binding to the target antigens. It is also important to note that these data showed that the reactivity of purified IgG in mild conditions or in whole bat sera did not significantly differ (**Fig. 4a**).

**Figure 4.**
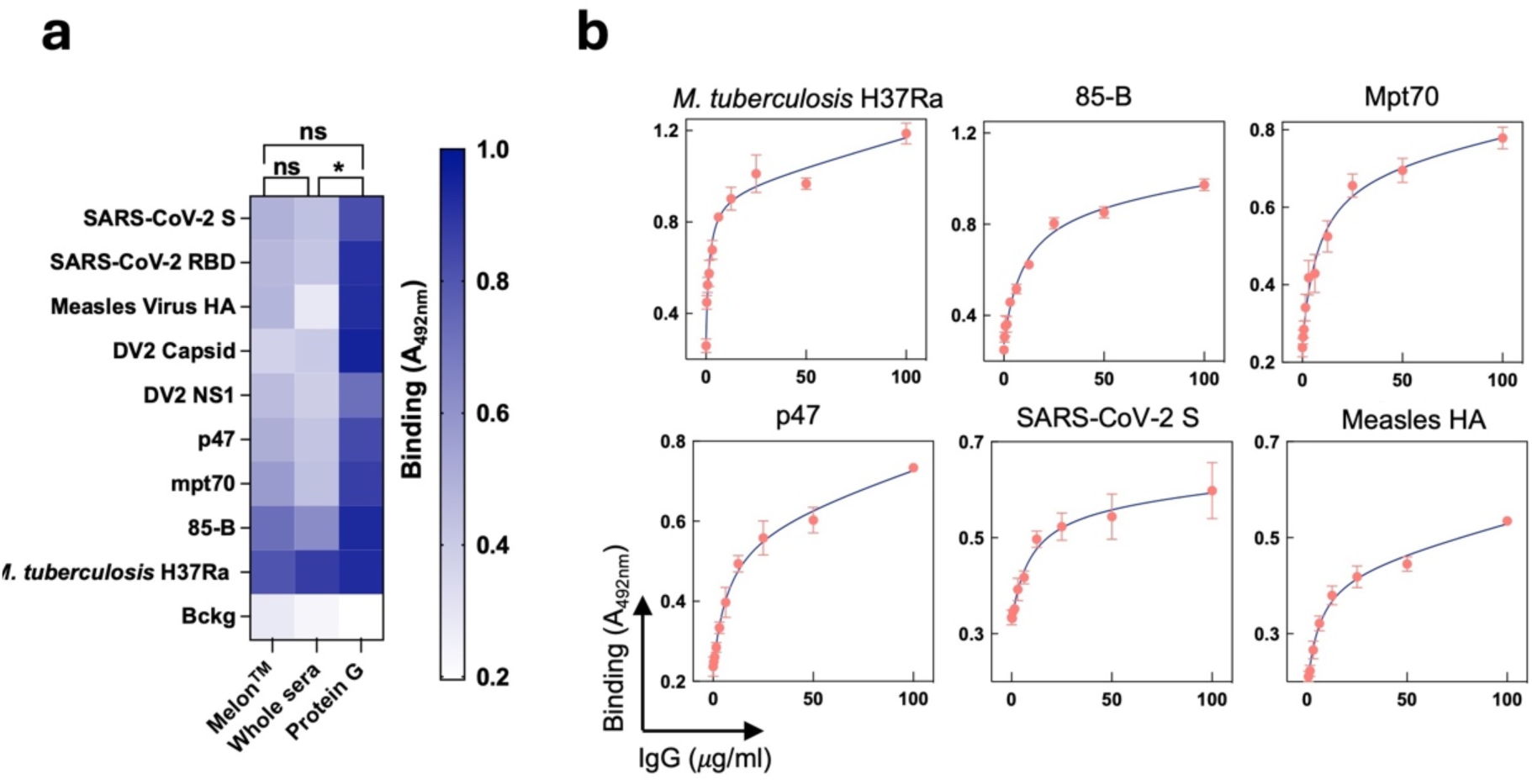
Binding of bat antibodies to intact bacteria or pathogen-derived proteins. **a**. The reactivity of bat antibodies purified by Melon^TM^, Protein G or in whole sera to pathogen-derived proteins. The heat map depicts the binding intensity. The values represented in the heat map are the averages of the absorbance readings (n=6) derived from two independent experiments. The statistical significance was evaluated using a one-way ANOVA multiple comparison Kruskal-Wallis test. The asterisk (*) denotes a P-value of 0.0032, while “ns” signifies non-significance. **b**. Concentration-dependent binding of IgG from *N. noctula* to the indicated pathogen-derived proteins or whole *M. tuberculosis* strain H37Ra bacteria. 85-B and Mpt70 derive from *M. tuberculosis,* and p47 from *T. pallidum*. Each point on the graph represents the average binding intensity (absorbance at 492 nm) ±SD from n=3, technical replicates.

### Analyses of pathogen specificities

The results presented herein demonstrate that the systemic serological assay proposed in the present study can be employed for the simultaneous detection of multiple pathogen-specificities of bat antibodies. It is postulated that most of the antibody reactivities to human pathogens identified in the present study represent cross-reactivity of antibodies against bat-specific pathogens that are closely related to human pathogens.

It can be inferred that the presence of antibodies in bats targeting pathogens such as SARS-CoV, *M. tuberculosis*, *M. leprae*, and P*. falciparum* can be attributed to the fact that bats are exposed to similar microorganisms with high sequence homology. Indeed, several bat species have been documented to harbor *Plasmodium sp*. and *Mycobacteria sp*. or phylogenetically related species, as well as diverse strains of coronaviruses, some of which exhibit close evolutionary relationships with SARS-CoV (13, 25–31). In light of these findings, sequence analyses of the SARS CoV epitopes that were significantly recognized by bat IgG in the present study revealed that the majority of these epitopes shared a high level or complete sequence identity with sequence stretches present in surface proteins from bat coronaviruses (**Suppl. Fig. 3**).

Furthermore, our findings indicate that bat antibodies recognize epitopes derived from certain human-specific pathogens that are unlikely to be present in bats, including HTLV-1, herpesvirus 4, dengue virus, and *Treponema pallidum*, among others (**Fig. 3 a,b**). The precise origin of these antibodies remains unknown. It is also conceivable that they may exhibit cross-reactivity to yet undescribed bat pathogens.

Finally, it seems plausible to suggest that some of the identified antigenic specificities of bat antibodies may have been generated following infection by the original species of pathogens. This group includes antibody specificities towards pathogens that can infect a broad range of mammals, such as *Borrelia burgdorferi*, *Staphylococcus aureus*, and *Toxoplasma gondii* (32–37)

Notwithstanding the potential for cross-reactivity in the use of human pathogen arrays, we contend that all antibody reactivities should be considered in the ongoing efforts to detect novel pathogens in bat populations. Furthermore, they could provide a foundation for targeted confirmatory genetic analyses.

### Limitations of the study

The primary limitation of the study is the use of an array comprising peptide epitopes derived from human pathogens, which may not fully reflect the complexity of the immune response in other species. It should be noted that some of the pathogens are known to infect only humans, and therefore may not be the most relevant targets for studying the immune responses in bats. Furthermore, the identified linear epitopes for arrays are selected for putative recognition by human B cells. Also, B cell epitopes derived from identical antigens can exhibit variation in bats. Furthermore, the array contains a disproportionately high number of epitopes derived from hepatitis C virus, comprising approximately 1,100 peptides out of a total of 4,300. This may introduce bias and lead to overestimation of antibody reactivity. Nevertheless, the array was applied here as a proof of concept, and its use as a readout does not compromise the overall approach presented in this study. Future studies should employ a tailored readout system, such as peptide or protein arrays, focused on antigens from bat-specific pathogens.

### Conclusive remarks

The present study proposes a rapid and effective experimental strategy for the analysis of bat IgG specificities at a systemic level. The method allows for the simultaneous analysis of antibody specificities towards a wide range of pathogens within a single day, which makes it suitable for monitoring wildlife diseases. Furthermore, the method does not require the use of bat-specific secondary antibodies, which allows for its broad applicability across different bat species and potentially other mammalian species. The illustration of extensive antigen reactivity within bat antibody repertoires will facilitate comprehensive analyses of pathogen dynamics at the ecosystem level. Furthermore, this approach can be invaluable for the study of anthropozoonoses, aiding in the protection of species and enhancing our understanding of disease transmission dynamics between humans and animals. The simplicity of the array’s customization allows for the creation of specific arrays tailored to a particular set of pathogens of concern. The comprehensive data output from these analyses may provide fundamental insight into bat-pathogen interactions.

## Methods

### Bat capture and sampling

Common noctule bats (*Nyctalus noctula*) were captured at Devetashka Cave, Bulgaria (N43.233, E24.885) in September 2023. Bats were trapped at cave entrances at sunset using mist nets and were subsequently placed individually in cotton bags to minimize stress and prevent injury. To collect blood samples, each bat was gently restrained, ensuring minimal stress. Blood was drawn from the uropatagium vein using a protocol that does not affect the survival rates of bats (38). A sterile 27-gauge insulin needle was used for venipuncture. Then, leaked blood was transferred directly into sterile microcentrifuge tubes using 20 µL sterile filter micropipette tips. Serum was obtained by blood centrifugation for 15-30 mins at 1000 × g and stored at -20℃ until use. The handling and sampling procedures were designed to minimize stress and potential harm to the bats. All activities were conducted in compliance with ethical guidelines and under the permit issued by the Bulgarian Biodiversity Act (No 927/04.04.2022). The researchers wore personal protective equipment during sampling. All non-disposable instruments and work surfaces were disinfected between handling each animal.

### Purification of IgG from bat sera

We compared the purification of IgG from pooled sera (n=35) of *N. noctula* using two approaches: 1) affinity chromatography with Protein G conjugated to an agarose matrix; 2) ion-exchange chromatography-based technology Melon^TM^ (Thermo Fischer Scientific, Waltham, MA). Both approaches are capable of purifying IgG from a diverse range of species across different mammalian orders.

The Protein G agarose matrix slurry (Thermo Fisher Scientific, Ref # 20397) was distributed in centrifugation spin columns with a volume of 0.2 ml per column. The matrix was washed a minimum of three times with sterile PBS (Gibco, Thermo Fisher Scientific, Ref # 14190144) by centrifugation for 1 min at 100 × g. Pooled bat sera (50 µL) was diluted 10 × in PBS at 0.5 ml of the solution and mixed with the settled matrix. The mixture was incubated for 20 min at 22 °C with end-to-end rotation. After four times washing with PBS, to the settled gel was added 0.5 ml of 0.1 M glycine pH 2.7. The mixture was incubated for 1 min, followed by collection of the elute by centrifugation (1 min at 100 × g). The eluates were immediately neutralized by adding 1/10 volume of Tris 1M, pH 9 and dialyzed against PBS.

The Melon^TM^ Gel IgG Spin Purification Kit (Thermo Fisher Scientific, Ref # 1859850) allows IgG purification in mild buffer conditions (pH 6.5-7). We followed the manufacturer’s protocol developed for the purification of human and mouse IgG with the sole exception that centrifugation steps were performed for 30 sec instead of 10 sec. The starting volume of pooled (n=35) bat sera was 50 µL.

The concentration of purified IgG from *N. noctula* was estimated by reading the absorbance at 280 nm using an absorbance spectrophotometer (NanoDrop, Thermo Fisher Scientific). The purity of IgG was assessed by SDS-PAGE performed in reducing and non-reducing conditions.

### Conjugation of bat IgG with fluorochrome

To label purified bat IgG with fluorochrome that has high quantum yield and stability, we used amine-reactive Alexa Fluor^TM^ 555 NHS ester (Thermo Fisher Scientific, Ref# A37571). The dry substance was dissolved in dimethyl sulfoxide (99.9 % purity, Sigma-Aldrich, St. Louis, MO, Ref# 472301) to a stock solution of 1.6 mM. The IgG was purified from N. noctula using the Melon™ system or Protein G and diluted in a buffer solution comprising two-thirds PBS and one-third 0.1M carbonate buffer (pH 9.3) to a final concentration of 2 µM. Subsequently, Alexa Fluor^TM^ 555 NHS was added, resulting in a final concentration of 40 µM. The reaction mixtures were incubated in the dark at 22 °C with continuous end-to-end rotation for one hour. The conjugation reaction was terminated by the addition of 10 mM Tris, resulting in a final pH of 8. Subsequently, to separate the fluorochrome-conjugated IgG from the unbound Alexa FluorTM 555 and to exchange the buffer, Sephadex G-25 resin-based (PD-10) columns were utilized (Cytiva, France Ref# 17085101). The columns were pre-equilibrated with PBS. The concentration and conjugation efficacy of IgG were estimated by absorbance spectrometry using an Agilent Cary-300 spectrophotometer (Agilent Technologies, Santa Clara, CA). The Alexa FluorTM 555-conjugated bat IgG was stored at 4 °C until required for use.

### Peptide microarray for assessment of bat IgG immunoreactivity

The PEPperCHIP® Infectious Disease Epitope Microarray (PEPperPRINT GmbH, Heidelberg, Germany) presents over 3,700 linear B-cell epitopes derived from 196 human pathogens. We consider these arrays an appropriate platform for providing proof of concept for high-throughput analyses of the specificity of bat antibodies. Due to the distinctive characteristics of the samples utilized in this study, substantial modifications to the manufacturer’s protocol were necessary.

To test the effect of the blocking agent, the microarray slides were alternatively incubated in PBS containing 0.05 % Tween 20 and 0.5 % BSA or 0.05 % Tween 20 and 1× Synthetic block (Invitrogen, Thermo Fisher Scientific, Ref# PA017). The slides were incubated for 30 min at 22 °C in 5 mL of blocking solutions using an orbital shaker at 50 rpm. After removing the blocking solutions, the arrays were washed with PBS containing 0.05 % Tween 20 (5 mL volume). Next, the microarray slides were incubated with IgG from *N. noctula* labeled with Alexa Fluor^TM^ 555 diluted to 5 µg/ml in PBS containing 0.05 % Tween 20. To compare the effect of the antibody purification method, microarrays were alternatively incubated with Protein G- or Melon^TM^ system-purified IgG. The arrays were incubated with 4 mL of antibody solution for 2 hours at 22 ° C. The incubation was performed on an orbital shaker at 50 rpm. Following incubation, the slides were washed by incubation 3× for one minute each with 5 mL of PBS containing 0.05 % Tween 20. The microarray slides were then soaked in deionized water to wash out the salts and dried under a stream of inert gas (argon). The immunoreactivity of bat IgG was estimated by reading the fluorescence intensity after excitation at 535 nm using a GenePix 4000B microarray scanner (Molecular Devices, San Jose, CA).

The data were analyzed using Spotxel software, version 1.7.7 (Sicasys, Heidelberg, Germany). The software enabled the quantification of mean fluorescence intensity, the identification of target peptides, and the calculation of background fluorescence for the array. Subsequent data analyses were performed using Microsoft Excel for Mac, version 16.86 (Microsoft, Redmond, WA), GraphPad Prism, version 10.2.3 (GraphPad Software, Boston, MA), and R Studio. The number of positive hits obtained for the binding of bat antibodies to epitopes from the hepatitis C virus was not displayed on the heat maps (Fig. 3a, b). The array contains approximately 1,100 peptides derived from the hepatitis C virus, representing a disproportionately high number compared to the total of approximately 4,300 peptides. This imbalance may introduce bias in the data, making comparisons with other pathogens irrelevant.

### ELISA

ELISA was performed using 384-well polystyrene plates (NUNC MaxiSorp, Thermo Fisher Scientific Ref# 442404). The plates were coated with recombinant SARS-CoV-2 Spike and SARS-CoV-2 RBD (both Wuhan strain, kindly provided by H. Mouquet, Institute Pasteur), as well as with the following proteins obtained from Immune Technology Corp. (New York, NY), Capsid Protein (Dengue virus 2, Ref# IT-006-015Ep), NS1 (Dengue virus 2, Ref# IT-006-0051p), Hemagglutinin Mosaic (Measles Virus, Ref# IT-026-002p), Antigen Mpt70 (M. Tuberculosis/H37Rv, Ref# IT-500-012Ep), Antigen 85-B (M. Tuberculosis/H37Rv, Ref# IT-500-015Ep), p47 (*T. pallidum*/Nichols strain, Ref# IT-501-004Ep). All proteins were diluted at 5 µg/ml in PBS. For coating with whole inactivated *Mycobacterium tuberculosis* H37Ra (BD, Ref# 231141), bacteria suspension was diluted to 100 µg/ml in PBS. The proteins or bacteria were incubated for 90 minutes at 22 °C. The plates were then blocked by incubation for one hour with a blocking solution consisting of PBS containing 1 × synthetic block and 0.25 % Tween 20. Purified IgG from *N. noctula* (intact, without conjugated fluorochrome) was diluted in PBS containing 0.05 % Tween 20 (PBS-T) to a fixed concentration of 20 µg/ml or first to 100 µg/ml, followed by two-fold dilutions in PBS-T down to 0.039 µg/ml. Alternatively, the reactivity of IgG in whole sera of *N. noctula* was also tested after 200 × dilution in PBS-T. The plates were incubated with antibodies for 90 minutes at 22 °C. After incubation, the plates were washed with PBS-T and further incubated with 2 µg/ml in PBS-T of biotinylated protein G (Pierce™, Thermo Fischer Scientific, Ref# 29988) for 1 hour at 22 °C. Following another washing step, streptavidin-HRP (Invitrogen, Thermo Fischer Scientific) diluted 100 × in PBS-T was added to the plates and incubated for 30 min at 22 °C. The immunoreactivity was developed by adding the substrate solution OPD SigmaFAST^TM^ (Sigma-Adrich, Ref# P9187). The reaction was terminated by adding 2M HCl. The absorbance was read at 492 nm with Infinite 200 Pro microplate reader (Tecan, Männedorf, Switzerland).

### Bioinformatics Analyses

Sequence analyses of epitopes recognized by bat antibodies with Z score 2 or greater were conducted using the Clustal Omega program (version 1.2.4) for multiple sequence alignment from EMBL’s European Bioinformatics Institute (https://www.ebi.ac.uk/). Sequence BLAST analyses were conducted using the BLASTP 2.16.0 algorithm. In the majority of cases, the UniProt online platform (https://www.uniprot.org/) was employed. In select instances, NCBI Protein Blast was utilized as an alternative (https://blast.ncbi.nlm.nih.gov/Blast.cgi).

## Acknowledgements

The experimental work was supported by a grant from the European Research Council (Project CoBABATI ERC-StG-678905), awarded to J.D.D. N.T. and V.Z. were supported by fellowships from the French Institute Bulgaria (Institut français de Bulgarie). The fieldwork for this study was supported by the Bulgarian National Science Fund, project КП-06-Н51/9 “Caves as a reservoir for novel and reoccurring zoonoses — ecological monitoring and metagenomic analysis in real-time.”

## Author contributions

N.T., V.Z., and J.D.D. conceived the study and designed the experiments; N.T., K.K., K.D., and V.Z. collected essential samples; N.T., F.E., R.V.L., M.L., and J.D.D. performed the experiments. A.P., and J.D.D. analyzed the obtained data; J.D.D. supervised the research; N.T. and J.D.D. wrote the manuscript; all authors contributed to editing of the manuscript.

