## Supplemental Figures for "An integrative approach for profiling antibody responses in bats to human pathogens"

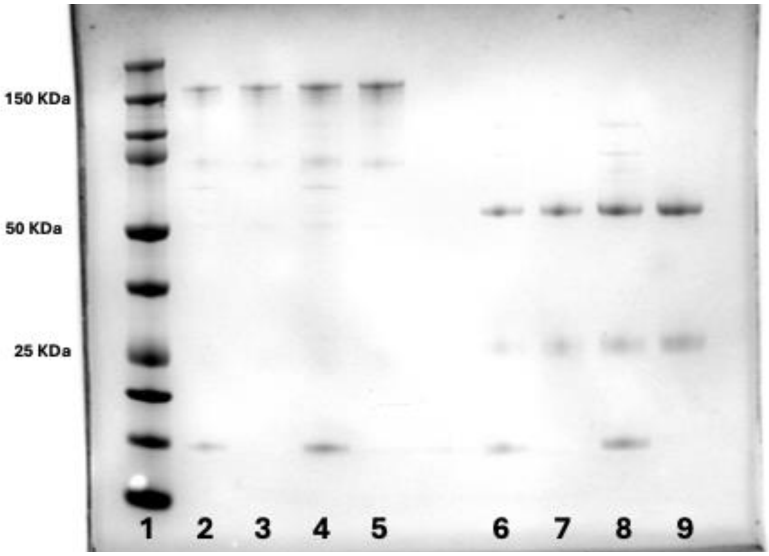


1. MW markers
2. IgG Melon Non-reducing 0.55 ug
3. IgG Protein G Non-reducing 0.55 ug
4. IgG Melon Non-reducing 1.11 ug
5. IgG Protein G Non-reducing 1.11 ug

6. IgG Melon Reducing 0.55 ug
7. IgG Protein G Reducing 0.55 ug
8. IgG Melon Reducing 1.11 ug
9. IgG Protein G Reducing 1.11 ug

**Supplemental Figure 1**. SDS-PAGE of IgG from *N. noctule.* The samples were tested in non-reducing and reducing conditions.

**Epitopes from *Staphylococcus aureus***
**ABC Transporter Atp-Binding Protein**

```
E5        ---------QPSSRRYPF  9
E1        ---------KVYVGNYDF  9
E4        --------IKVYVGNYD-  9
E3        -----TFLRGFLGR---   9
E2        KWGVTTSLS--------   9
E6        ---KTTLLKI-LA----   9
```

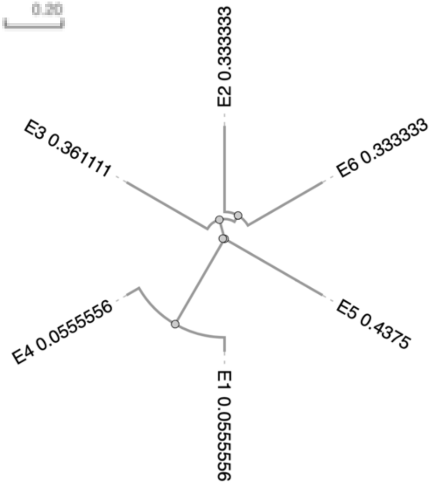


**Epitopes from Dengue virus 2**
**Envelope Glycoprotein Dengue**

```
E5        ------------MAKNKP 6
E2        ---------CRLRMD---- 6
E6        -----GVIITW------- 6
E4        AELTGYGTVT-------- 10
E1        IGISNRDFV--------- 9
E7        --VSGGSWVDIVLE---- 12
E3        ---------LDFELI---- 6
```

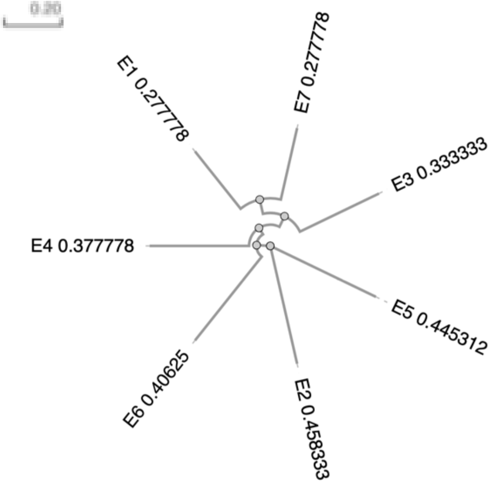


***Orientia tsutsugamushi***
**60 kDa Chaperonin**

```
E2        ---DRGYISQY--    8
E5        -----GYISQYFA    8
E1        EDNNTGNR-----    8
E3        -DKNNSDIE----    8
E4        -DSKNFNFE----    8
```

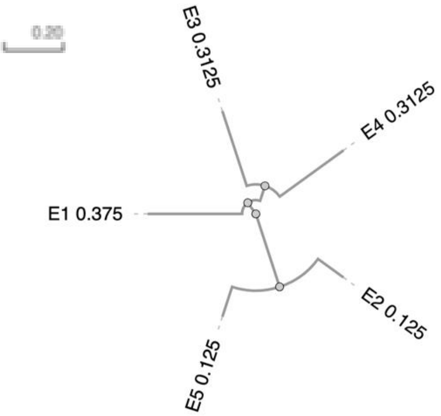


**Hepatitis Delta virus**
**Delta Antigen**

```
E4        ----DILFPADPPFSPQSC-    15
E5        -----------PPFSPQ---    6
E3        ---------KDGEGAPPAKKL   12
E1        ----------GSQGFP-----   6
E2        ENKKKQLSAGGKNLS-----    15
```

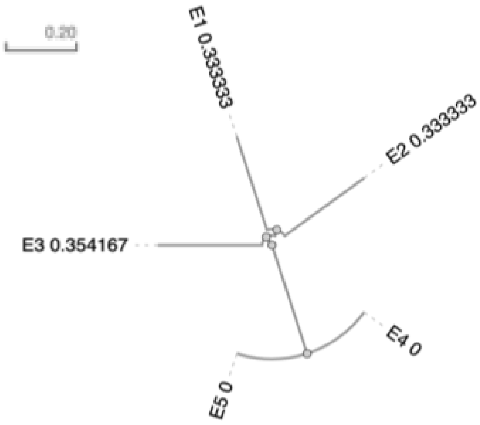


**Supplemental Figure 2**. Sequence alignment of epitopes from epitopes derived from the indicated pathogen proteins that are recognized with Z scores 2 or greater. The red color is used to highlight the identical motives. The sequence alignment plot and phylogenetic analyses (Guide trees) were generated by the Clustal Omega program (version 1.2.4) for multiple sequence alignment (https://www.ebi.ac.uk/).

| Epitope E1 | | % Identity |
|---|---|---|
| SARS CoV-1 | PSSKRFQPFQQFGRD | 100 |
| Bat CoV HKU3 | -SSKRFSPFQQFGRD | 87 |
| Bat CoV Rp3/2004 | -SSKRFQSFQQFGRD | 87 |
| Bat CoV 279 | -SSKRFQSFQQFGRD | 87 |
| Bat CoV Cp/Yunnan2011 | -SSKKFQSFQQFGRD | 80 |
| Bat SARS-like CoV YNLF_31C | -SSKTFQSFQQFGRD | 80 |
| SARS CoV-2 | -SNKKFLPFQQFGRD | 73 |

| Epitope E2 | | % Identity |
|---|---|---|
| SARS CoV-1 | HEGKAYFPREGV | 100 |
| Bat CoV HKU3 | HEGKAYFPREGV | 100 |
| Bat CoV Rp3/2004 | HEGKAYFPREGV | 100 |
| Bat CoV 279 | HEGKAYFPREGV | 100 |
| Bat SARS-like CoV YNLF_31C | HEGKAYFPREGV | 100 |
| Bat CoV Cp/Yunnan2011 | HQGKAYFPREGV | 92 |
| SARS CoV-2 | HDGKAHFPREGV | 83 |

| Epitope E3 | | % Identity |
|---|---|---|
| SARS CoV-1 | AGCLIGAEHVDTSYE | 100 |
| Bat SARS-like CoV YNLF_31C | AGCLIGAEHVNSSYE | 87 |
| Bat CoV HKU3 | AGCLIGAEHVNASYE | 87 |
| Bat CoV Rp3/2004 | AGCLIGAEHVNASYE | 87 |
| Bat CoV 279 | AGCLIGAEHVNASYE | 87 |
| Bat CoV Cp/Yunnan2011 | AGCLIGAEHVNASYE | 87 |
| SARS CoV-2 | AGCLIGAEHVNNSYE | 87 |

| Epitope E4 | | % Identity |
|---|---|---|
| SARS CoV-1 | LCPFGEVFNATKFPS | 100 |
| SARS CoV-2 | LCPFGEVFNATRFAS | 87 |
| Bat SARS-like CoV YNLF_31C | LCPFDKVFNATRFPS | 80 |
| Bat CoV HKU3 | -CPFDKVFNATRFP- | 67 |
| Bat CoV Rp3/2004 | -CPFDKVFNATRFP- | 67 |
| Bat CoV Cp/Yunnan2011 | -CPFDRVFNASRFPS | 67 |
| Bat CoV 279 | -CPFDKVFNASRFP- | 60 |

| Epitope E5 | | % Identity |
|---|---|---|
| SARS CoV-1 | DLPSGFNTLKPIFKL | 100 |
| SARS CoV-2 | DLPSGFNTLKPIFKL | 100 |
| Bat CoV HKU3 | -LPTGFSVLKPILKL | 67 |
| Bat SARS-like CoV YNLF_31C | -LPAGLSVLKPILKL | 60 |
| Bat CoV Rp3/2004 | -LPIGFSVLRPILKL | 60 |
| Bat CoV Cp/Yunnan2011 | --PVGFSVLKPILKL | 60 |
| Bat CoV 279 | --PAGFSVLRPILKL | 53 |

| Epitope E6 | | % Identity |
|---|---|---|
| SARS CoV-1 | DLGDISGINASVVNI | 100 |
| SARS CoV-2 | DLGDISGINASVVNI | 100 |
| Bat CoV HKU3 | DLGDISGINASVVNI | 100 |
| Bat CoV Rp3/2004 | DLGDISGINASVVNI | 100 |
| Bat CoV 279 | DLGDISGINASVVNI | 100 |
| Bat SARS-like CoV YNLF_31C | DLGDISGINASVVNI | 100 |
| Bat CoV Cp/Yunnan2011 | DLGDISGINASVVNI | 100 |

| Epitope E7 | | % Identity |
|---|---|---|
| SARS CoV-1 | PFGEVFNATKFPSVY | 100 |
| SARS CoV-2 | PFGEVFNATRFASVY | 87 |
| Bat SARS-like CoV YNLF_31C | PFDKVFNATRFPSVY | 80 |
| Bat CoV HKU3 | PFDKVFNATRFPNVY | 73 |
| Bat CoV Rp3/2004 | PFDKVFNATRFPNVY | 73 |
| Bat CoV Cp/Yunnan2011 | PFDRVFNASRFPSVY | 73 |
| Bat CoV 279 | PFDKVFNASRFPNVY | 67 |

| Epitope E8 | | % Identity |
|---|---|---|
| SARS CoV-1 | QLIRAAEIRASANLA | 100 |
| SARS CoV-2 | QLIRAAEIRASANLA | 100 |
| Bat CoV HKU3 | QLIRAAEIRASANLA | 100 |
| Bat CoV Rp3/2004 | QLIRAAEIRASANLA | 100 |
| Bat CoV 279 | QLIRAAEIRASANLA | 100 |
| Bat SARS-like CoV YNLF_31C | QLIRAAEIRASANLA | 100 |
| Bat CoV Cp/Yunnan2011 | QLIRAAEIRASANLA | 100 |

| Epitope E9 | | % Identity |
|---|---|---|
| SARS CoV-1 | KNQCVNFNFNGLTGT | 100 |
| SARS CoV-2 | KNKCVNFNFNGLTGT | 93 |
| Bat CoV HKU3 | KNQCVNFNFNGLKGT | 93 |
| Bat CoV Rp3/2004 | KNQCVNFNFNGLKGT | 93 |
| Bat CoV 279 | KNQCVNFNFNGLRGT | 93 |
| Bat CoV Cp/Yunnan2011 | KNQCVNFNFNGLKGT | 93 |
| Bat SARS-like CoV YNLF_31C | KNQCVNFNFNGFKGT | 87 |

| Epitope E10 | | % Identity |
|---|---|---|
| SARS CoV-1 | PSGFNTLKPIFKLPL | 100 |
| SARS CoV-2 | PSGFNTLKPIFKLPL | 100 |
| Bat CoV HKU3 | PTGFSVLKPILKLP- | 67 |
| Bat CoV Cp/Yunnan2011 | PVGFSVLKPILKLP- | 67 |
| Bat CoV Rp3/2004 | PIGFSVLRPILKLP- | 60 |
| Bat CoV 279 | PAGFSVLRPILKLP- | 60 |
| Bat SARS-like CoV YNLF_31C | PAGLSVLKPILKLP- | 60 |

| Epitope E11 | | % Identity |
|---|---|---|
| SARS CoV-1 | SLQTYVTQQLIRAAE | 100 |
| SARS CoV-2 | SLQTYVTQQLIRAAE | 100 |
| Bat CoV HKU3 | SLQTYVTQQLIRAAE | 100 |
| Bat CoV Cp/Yunnan2011 | SLQTYVTQQLIRAAE | 100 |
| Bat CoV Rp3/2004 | SLQTYVTQQLIRAAE | 100 |
| Bat CoV 279 | SLQTYVTQQLIRAAE | 100 |
| Bat SARS-like CoV YNLF_31C | SLQTYVTQQLIRAAE | 100 |

| Epitope E12 | | % Identity |
|---|---|---|
| SARS CoV-1 | PTNFSISITTEVMPV | 100 |
| Bat CoV HKU3 | PTNFSISVTTEVMPV | 93 |
| Bat CoV Cp/Yunnan2011 | PTNFSISVTTEVMPV | 93 |
| Bat CoV Rp3/2004 | PTNFSISVTTEVMPV | 93 |
| Bat CoV 279 | PTNFSISVTTEVMPV | 93 |
| Bat SARS-like CoV YNLF_31C | PTNFSISVTTEVMPV | 93 |
| SARS CoV-2 | PTNFTISVTTEILPV | 73 |

| Epitope E13 | | % Identity |
|---|---|---|
| SARS CoV-1 | PKTSEILDISPCAFG | 100 |
| Bat CoV HKU3 | PQTLEILDISPCSFG | 80 |
| Bat CoV Rp3/2004 | PQTLEILDISPCSFG | 80 |
| Bat CoV 279 | PQTLEILDISPCSFG | 80 |
| Bat SARS-like CoV YNLF_31C | PKTLQILDISPCSFG | 80 |
| Bat CoV Cp/Yunnan2011 | PQTLQILDISPCSFG | 73 |
| SARS CoV-2 | PQTLEILDITPCSFG | 73 |

| Epitope E14 | | % Identity |
|---|---|---|
| SARS CoV-1 | SPDGKPCTPPALNCY | 100 |
| SARS CoV-2 | SPDGKPCTPPALNCY | 100 |
| Bat CoV HKU3 | | <50 |
| Bat CoV Rp3/2004 | | <50 |
| Bat CoV 279 | | <50 |
| Bat SARS-like CoV YNLF_31C | | <50 |
| Bat CoV Cp/Yunnan2011 | | <50 |

| Epitope E15 | | % Identity |
|---|---|---|
| SARS CoV-1 | AQDIWGTSAAAYFVG | 100 |
| SARS CoV-2 | AQDIWGTSAAAYFVG | 100 |
| Bat CoV HKU3 | | <50 |
| Bat CoV Rp3/2004 | | <50 |
| Bat CoV 279 | | <50 |
| Bat SARS-like CoV YNLF_31C | | <50 |
| Bat CoV Cp/Yunnan2011 | | <50 |

| Epitope E16 | | % Identity |
|---|---|---|
| SARS CoV-1 | GDISGINASVVNIQK | 100 |
| SARS CoV-2 | GDISGINASVVNIQK | 100 |
| Bat CoV HKU3 | GDISGINASVVNIQK | 100 |
| Bat CoV Rp3/2004 | GDISGINASVVNIQK | 100 |
| Bat CoV 279 | GDISGINASVVNIQK | 100 |
| Bat SARS-like CoV YNLF_31C | GDISGINASVVNIQK | 100 |
| Bat CoV Cp/Yunnan2011 | GDISGINASVVNIQK | 100 |

| Epitope E17 | | % Identity |
|---|---|---|
| SARS CoV-1 | IVAYTMSLGADSSIA | 100 |
| Bat CoV HKU3 | IVAYTMSLGAENSIA | 87 |
| Bat CoV Rp3/2004 | IVAYTMSLGAENSIA | 87 |
| Bat CoV 279 | IVAYTMSLGAENSIA | 87 |
| Bat CoV Cp/Yunnan2011 | IVAYTMSLGAENSIA | 87 |
| Bat SARS-like CoV YNLF_31C | IVAYTMSLGAENSVA | 80 |
| SARS CoV-2 | IIAYTMSLGAENSVA | 73 |

| Epitope E18 | | % Identity |
|---|---|---|
| SARS CoV-1 | YRFNGIGVTQNVLYE | 100 |
| SARS CoV-2 | YRFNGIGVTQNVLYE | 100 |
| Bat CoV HKU3 | YRFNGIGVTQNVLYE | 100 |
| Bat CoV Rp3/2004 | YRFNGIGVTQNVLYE | 100 |
| Bat CoV 279 | YRFNGIGVTQNVLYE | 100 |
| Bat CoV Cp/Yunnan2011 | YRFNGIGVTQNVLYE | 100 |
| Bat SARS-like CoV YNLF_31C | YRFNGIGVTQNVLYE | 100 |

| Epitope E19 | | % Identity |
|---|---|---|
| SARS CoV-1 | HTINHTFGNPVIPFK | 100 |
| SARS CoV-2 | HTINHTFGNPVIPFK | 100 |
| Bat CoV HKU3 | | <50 |
| Bat CoV Rp3/2004 | | <50 |
| Bat CoV 279 | | <50 |
| Bat CoV Cp/Yunnan2011 | | <50 |
| Bat SARS-like CoV YNLF_31C | | <50 |

| Epitope E20 | | % Identity |
|---|---|---|
| SARS CoV-1 | IFLLFLTLTSGSDLD | 100 |
| SARS CoV-2 | IFLLFLTLTSGSDLD | 100 |
| Bat CoV HKU3 | | <50 |
| Bat CoV Rp3/2004 | | <50 |
| Bat CoV 279 | | <50 |
| Bat CoV Cp/Yunnan2011 | | <50 |
| Bat SARS-like CoV YNLF_31C | | <50 |

**Supplemental Figure 3**. Sequence identity of epitopes from SARS CoV recognized by bat IgG with sequence from closely related coronaviruses. Sequences of epitopes from SARS CoV E2 glycoprotein (top row) were aligned with sequences from coronaviruses with highest identity. The lack of identity as compared to the original sequence was indicated in red. The assessment of sequence identities was done by UniProt (https://www.uniprot.org/) or NCBI Protein Blast (https://blast.ncbi.nlm.nih.gov) online platforms.
